# Idiosyncratic choice bias and feedback-induced bias differ in their long-term dynamics

**DOI:** 10.1101/2024.09.02.610741

**Authors:** Lior Lebovich, Lea Kaplan, David Hansel, Yonatan Loewenstein

**Affiliations:** The Edmond and Lily Safra Center for Brain Sciences, The Hebrew University, Jerusalem, Israel; Department of Collective Behaviour, Max Planck Institute of Animal Behavior, Konstanz, Germany; Centre for the Advanced Study of Collective Behaviour, University of Konstanz, Universitätsstraße 10, 78464, Konstanz, Germany; Cerebral Dynamics, Plasticity and Learning Laboratory, CNRS, 45 Rue des Saints Pères, 75270, Paris, France; The Alexander Silberman Institute of Life Sciences, The Hebrew University, Jerusalem, Israel; Dept. of Cognitive Sciences and The Federmann Center for the Study of Rationality, The Hebrew University, Jerusalem, Israel

## Abstract

A well-known observation in repeated-choice experiments is that a tendency to prefer one response over the others emerges if the feedback consistently favors that response. Choice bias, a tendency to prefer one response over the others, however, is not restricted to biased-feedback settings and is also observed when the feedback is unbiased. In fact, participant-specific choice bias, known as idiosyncratic choice bias (ICB), is common even in symmetrical experimental settings in which feedback is completely absent. Here we ask whether feedback-induced bias and ICB share a common mechanism. Specifically, we ask whether ICBs reflect idiosyncrasies in choice-feedback associations *prior* to the measurement of the ICB. To address this question, we compare the long-term dynamics of ICBs with feedback-induced biases. We show that while feedback effectively induced choice preferences, its effect is transient and diminished within several weeks. By contrast, we show that ICBs remained stable for at least 22 months. These results indicate that different mechanisms underlie the idiosyncratic and feedback-induced biases.

## Introduction

Perceptual decision making is often studied using the 2-alternative forced choice (2AFC) paradigm, in which the experimentalist presents two stimuli that vary along one physical dimension, and the participant is instructed to choose the ‘larger’ one. In many of these experiments, participants receive feedback after every trial, which can bias their choices. For example, if the feedback indicates that choosing one of the alternatives was the correct response, participants are more likely to choose it again in the following trial. This tendency can be viewed as a form of operant learning, in which participants bias their choices in a direction that they deem more likely to be rewarded. If over many trials, the feedback is asymmetric, consistently favoring one alternative over the other, participants will develop a substantial preference in favor of that alternative ^1–3^.

Somewhat surprisingly, however, participants exhibit preferences in 2AFC experiments, even when the feedback is unbiased. These preferences are idiosyncratic: different participants exhibit a preference in favor of different alternatives and the magnitudes of these preferences vary across participants^4–13^. These idiosyncratic choice biases (ICBs) are observed even in well-controlled experiments and in absence of feedback^14^. Finally, ICBs are not restricted to humans. They are also observed in animals ranging from flies^15^ to rodents^16–20^ to monkeys^21,22^.

In a previous study^14^ we hypothesized that ICBs are the result of idiosyncratic microscopic heterogeneities in connectivity of the neural circuits involved in the decision-making process. However, an appealing alternative hypothesis is that ICBs reflect idiosyncratic feedback that the participants received *prior* to the experiment. Participants are not *tabula rasa* when they start the experiment, and their life-experience involves idiosyncratic interactions with the environment followed by feedback. According to this hypothesis, ICBs reflect idiosyncratic feedback.

Clearly, the latter hypothesis cannot be addressed directly, because we have no access to the full life-history of the participants, and even if we had, it would be very difficult to determine which of the life experiences is *relevant* to any 2AFC task. Here we adopt an indirect approach: If ICBs reflect an idiosyncratic history of feedback-induced choice bias, then we expect that they will be as stable (or unstable) as biases induced in the laboratory using feedback. We report here that this is not the case. We find that while choice bias can be readily induced by feedback, it is relatively short-lived, decaying within weeks after its induction. By contrast, ICBs are remarkably stable, hardly changing in a 22 months period. These results indicate that ICBs are not the result of idiosyncratic feedback, consistent with the hypothesis that they result from irreducible heterogeneities in brain connectivity.

## Results

### Characterization of the ICBs

We used the vertical bisection discrimination task, depicted in Fig. 1A (inset) to study ICBs (Methods). In each trial, a vertical transected line was presented on the screen and participants were instructed to indicate which of the two vertical segments was longer. No trial-by-trial feedback was given to the participants (*n* = 183) about their answers. We computed the fraction of responses in which the participants reported that the upper segment of the transected line was longer than the bottom segment, a quantity which we denote by *p*_u_, separately for each participant and offset. Figure 1A depicts *p*_u_ for three participants as a function of the offset. As expected, *p*_u_ increased with the offset, 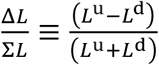,where *L*^u^ and *L*^d^ denote the lengths of the up and down segments of the vertical line. Notably, the participant denoted by a blue line was biased in favor of responding ‘down’ even in the trials in which Δ*L* was positive (but small). By contrast, the psychometric curve of the participant denoted by the red curve is shifted to the left relative to the blue curve, indicating a bias in favor of responding ‘up’. To quantify the idiosyncratic preferences towards and against responding ‘up’, we focused on the choices of the participants in ‘impossible’ trials, in which the line was transected at its veridical midpoint (Δ*L* = 0, 40/240 trials). We denote by 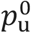 the fraction of ‘up’ responses of a participant in these trials. While the participant whose psychometric curve is in black in Fig. 1A did not exhibit any significant bias (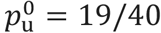trials, *p* = 0.87, two-sided Binomial test), the participants whose psychometric curves are plotted in red and blue were significantly biased (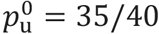and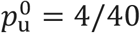, respectively; *p*<0.001, two-sided Binomial tests).

**Fig. 1.**
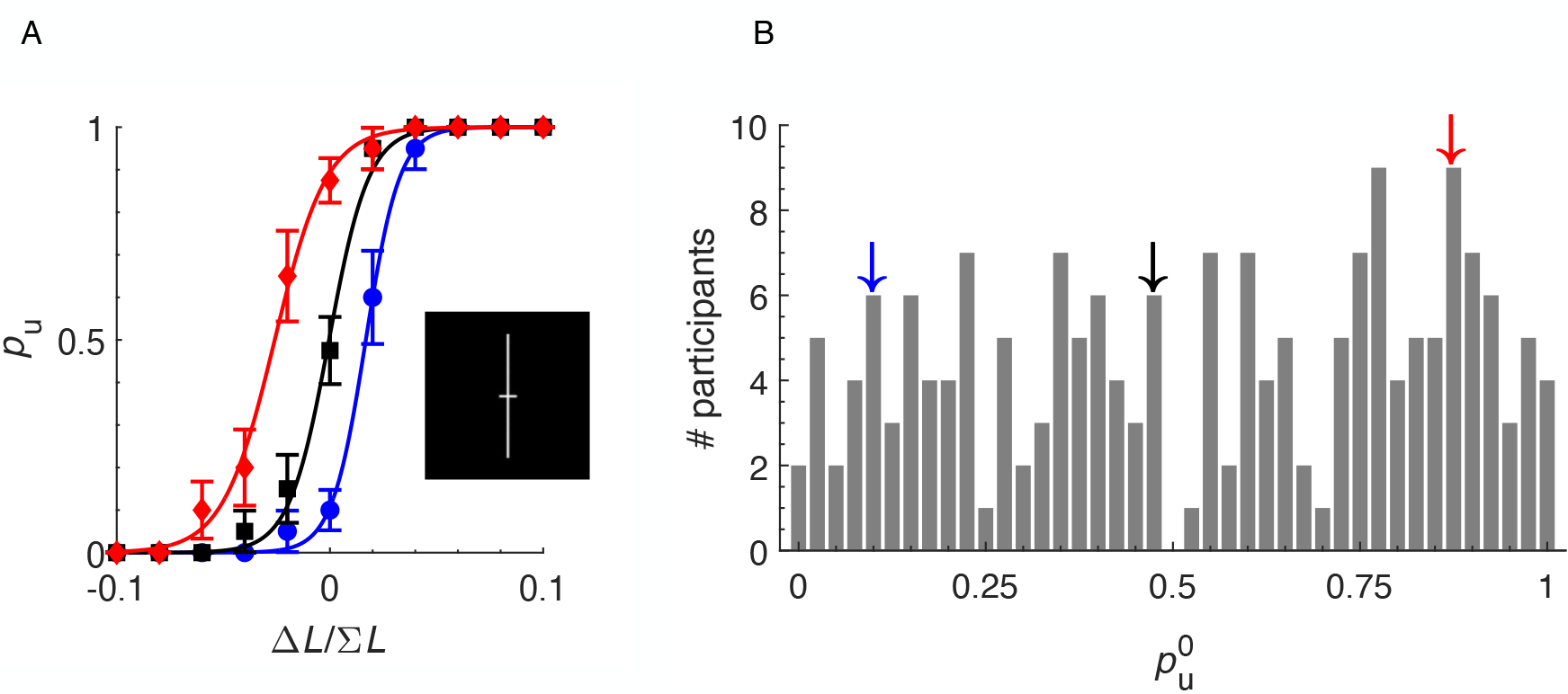
ICBs in the vertical bisection task. **A**, psychometric curves of three participants. The observed fraction, *p*_U_, of responding ‘up’, is plotted as a function of the sensory offset, 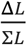. Error bars denote the standard error of the mean (SEM). Solid lines are best-fit logistic functions. One participant did not exhibit any significant bias (black), one was biased in favor of responding up (blue) and one (red) against it. Inset: the stimulus in a single trial. **B**, the distribution across the participants (*n* = 183) of 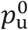, the value of *p*, in trials in which Δ*L* = 0. Arrows denote 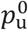 for the three participants in panel A, color coded.

The distribution of 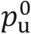 is depicted in Fig. 1B. Across the participants, 69% exhibited a significant bias (39% with 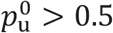 and 30% with 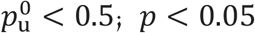, two-sided Binomial tests, not corrected for multiple comparisons). The mean of the distribution, 0.53, is not significantly different from chance (95% CI 0.49-0.58, bootstrap). The variance of the distribution was significantly larger than expected by chance (*p* < 0.001, two-sided bootstrap test, Bernoulli process; *n* = 183 participants, 40 trials per participant) further establishing the existence of ICBs in the task. Finally, the ICBs were not restricted to the impossible trials. Rather, they manifested as idiosyncratic lateral shifts of the psychometric curve. Indeed, across the participants, the probability of choosing ‘up’ in ‘possible trials’ was highly correlated with that probability in the impossible trials (Supplementary Fig. S1; two-sided Pearson’s ρ = 0.82, *p* < 0.001).

### ICBs are stable over many months

To study how ICBs change over time, the ICB of each of the participants depicted in Fig. 1B was measured again later (session 2 in Fig. 2A; also see: Methods). One subgroup of 29 participants was also tested 22 months after the second session (session 3).

**Fig. 2.**
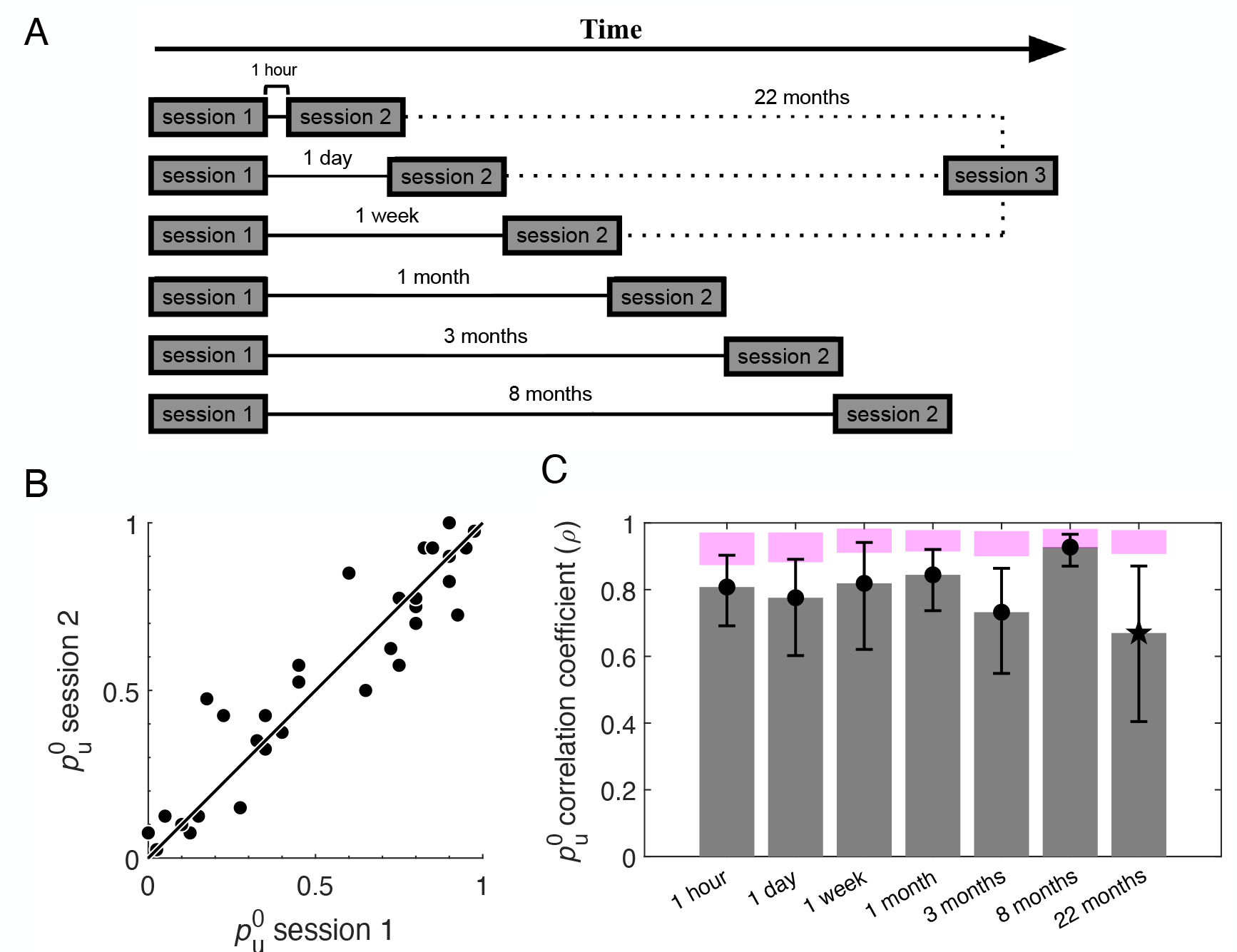
ICB is stable over months. **A**, The experimental design. The ICBs of *n* = 183 participants were measured in the first session (session 1). Subsets of the participants were tested again in an identical session (session 2) 1 hour (*n* = 29), 1 day (*n* = 32), 1 week (*n* = 27), 1 month (*n* = 33), 3 months (*n* = 30) or 8 months later (*n* = 32). A subset (*n* = 29) also participated in a third session (session 3), 22 months after session 2 (dashed lines). **B**, ICB is stable over 8 months. Dots: participants tested in session 2, 8 months after session 1. The diagonal is plotted for comparison. **C**, ICBs are stable over time. Gray bars: Pearson correlation of 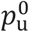 between session 1 and 2 (circles) and between session 2 and session 3 (star). Error bars: 95% confidence interval, bootstrap. Magenta area: 95% confidence interval expected under complete stability, bootstrap Bernoulli processes.

Figure 2B, plots 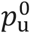 in session 2 *vs*. its value 8 months later (32 participants, Supplementary Fig. S2 for all other groups). It shows that 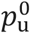 hardly changed during that delay. Quantifying this change, the correlation between the values of 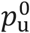 in the two sessions 8 months apart, was ρ = 0.82, which is not smaller than the correlation when the two sessions were 3 months, 1 month, 1 week, 1 day or even 1 hour apart (Fig. 2C). Even the correlation between the values of 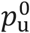 measured when the two sessions were 22 months apart (ρ = 0.67) was not significantly smaller than when the interval was 1 hour, 1 day, 1 week, 1 month or 3 months (*p* > 0.05. However, it was significantly smaller than for an interval of 8 months (*p* = 0.001, two-sided z-tests for the difference in correlation after Fisher z-transformation, not corrected for multiple comparisons). We therefore concluded that the participants retained their ICB even after many months.

### The dynamics of feedback-induced bias

If the observed ICBs reflect idiosyncratic histories of operant learning, we expect comparable stability for choice biases induced by providing proper feedback to the participants. Therefore, we set out to study the dynamics of such feedback-induced effect. To that goal, we recruited a group of *n* = 136 new participants and tested each one in three experimental sessions (Fig. 3). In session 1 we measured their ICBs without providing trial-by-trial feedback. This was not the case in the second session (session 2), 1 day later, where feedback was provided. In the session 3, either 1 day later or 1 month later, we again measured their ICBs without trial-by-trial feedback.

**Fig. 3.**
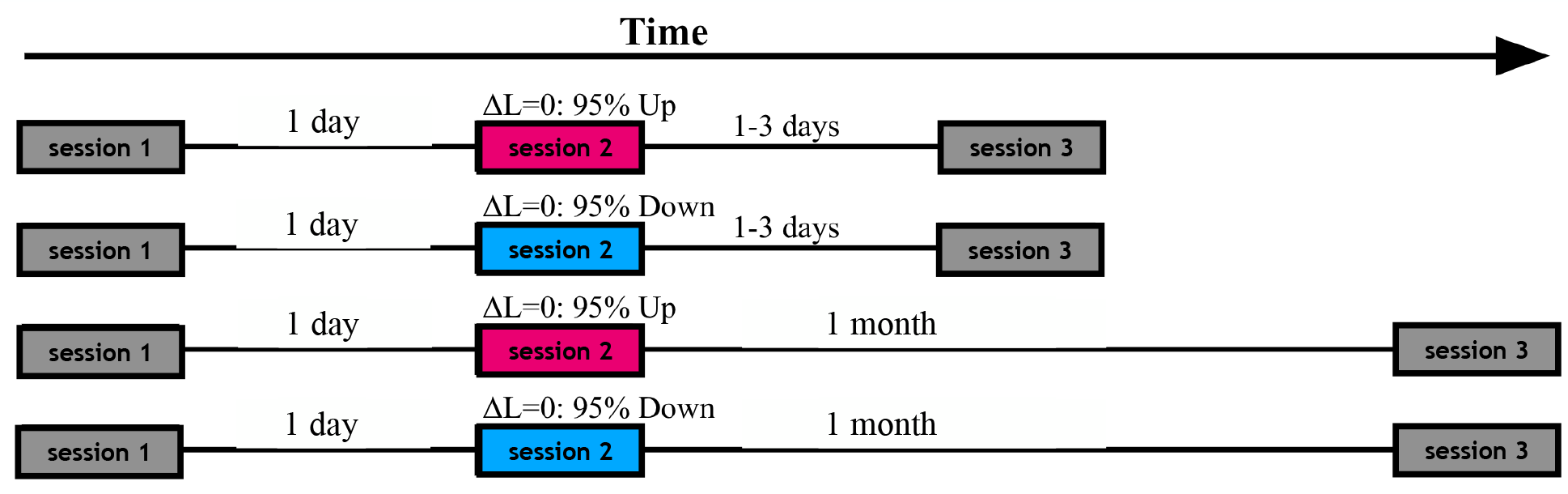
The experimental design to investigate the dynamics of feedback-induced bias. All participants (*n* = 136) participated in three sessions. In sessions 1 and 3 (gray) no trial-to-trial feedback was provided. In session 2, choices in all trials were followed by feedback. Feedback in 200 possible trials was accurate. Impossible trials (40 trials) were followed by asymmetrical feedback about the ‘correct’ response. Pink: ‘enhance up’ condition. The feedback indicated that the ‘correct’ response is ‘up’ in 38 impossible trials and ‘down’ in 2 impossible trials. Blue: ‘enhance down’ condition. The feedback indicated that the ‘correct’ response is ‘up’ in 2 impossible trials and ‘down’ in 38 impossible trials. All participants performed session 2 one day after their completion of session 1. Half of them participated in session 3 one day after completing session 2 (top) and the other half participated in session 3, 1 month after completing session 2 (bottom). Participants were divided into the 4 groups (2 feedback x 2 delay) by matching their ICBs in session 1 (Methods and Supplementary Fig. S3B).

As expected, since session 1 was identical to the first session of Fig. 2A, the distribution of ICBs over the participants was comparable in these two situations (Supplementary Fig. S3A). While in session 2, participants were presented with the same stimuli and the same task as in session 1, they also received, in every trial, feedback indicating whether their choice was correct (happy emoji) or incorrect (sad emoji). While the feedback was accurate in all possible trials (Δ*L* ≠ 0; 200/240 of the trials), it was biased in the impossible trials (Δ*L* = 0; 40/240 of the trials), to systematically influence the participants’ preferences. Specifically, participants were divided into two groups with different feedback schedules. In one group, ‘up’ response was considered ‘correct’ in 95% of the impossible trials while ‘down’ response was considered ‘correct’ in 5% of the trials. We refer to this group of participants as ‘enhance up’. In the second group, ‘enhance down’, the feedback was biased in the opposite direction: ‘up’ response was considered ‘correct’ in 5% of the impossible trials and ‘down’ response was correct in the other 95%.

So far, we separately considered each participant and averaged their responses over all the trials. To study how the feedback in session 2 changed the tendency to choose ‘up’ over trials, we separately considered the impossible trials and averaged the responses over the participants. To clarify the difference between averaging over responses given a participant and averaging over participants given a trial, we denote by 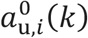 the response of participant *i* in impossible trial 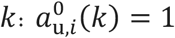 if participant *i* chose ‘up’ in impossible trial *k* (*k* ∈ {1, …, 40}), and 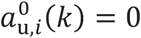 otherwise. The fraction of impossible trials in a session in which a participant *i* chose ‘up’ is 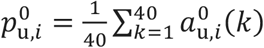. Note that the quantity 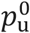 considered so far for each participant is in fact 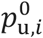, where the index *i* was omitted for simplicity of notations.

To study how feedback changed the tendency of choosing ‘up’ over trials, we averaged over participants, rather than over trials. We computed 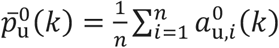, where *n* denotes the number of participants in each “enhance” group. The red and blue lines in Figure 4A depict a running average of 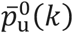 for the ‘enhance up’ and the ‘enhance down’ groups, respectively. Initially, the two groups exhibited comparable population averages. However, with trials, the effect of the feedback became more pronounced. These results are consistent with previous studies that demonstrated that feedback in even a small number of impossible trials can substantially bias participants’ choices^1^.

**Fig. 4.**
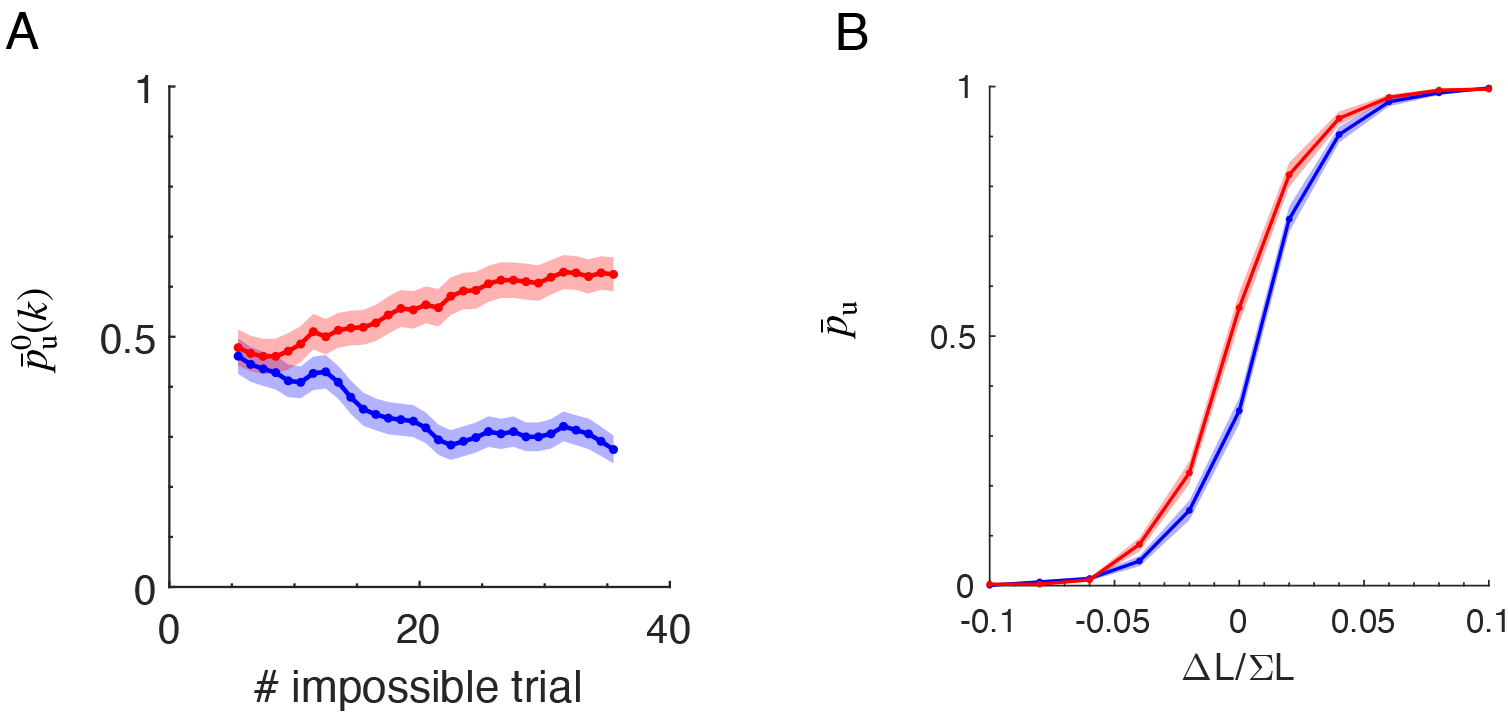
Feedback biases choices in session 2. **A**, the participant-averaged bias in the impossible trials,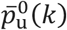, smoothed with a sliding window of 10 trials. Red and Blue: ‘enhance up’ and ‘enhance down’ groups of participants. Shaded area: SEM. **B**, The average (over participants) of the psychometric curve 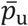 and SEM (shaded area) as a function of the relative difference between the length of the segments,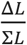.Colors as in **A**.

In our experiment, the biased feedback (95%; 5%) was applied only to the impossible trials, while it was correct and unbiased (50%; 50%) in the possible trials. A previous study has argued that the effect of biased feedback is selective to the specific stimuli that were associated with that feedback^23^. On the other hand, in a previous study that utilized auditory stimuli we showed that feedback in impossible trials also affects those which are possible ^1^. To test whether the feedback also affects choices in the possible trials in our bisection task, we compared the average psychometric curve of the ‘enhance up’ (red) and ‘enhance down’ (blue) participants in the second session (Fig. 4B). We found that the biased feedback shifted the entire psychometric curve, indicating that it also affected the possible trials.

If ICBs are the result of idiosyncratic feedback prior to the experiment, we predict the feedback-induced bias and ICBs to be equally stable. We therefore asked: how long would the feedback-induced bias in sessions 2 affect the bias of the participants? To address this question, half of the participants (68/136) were tested 1 day after the manipulation whereas half of the participants were tested after 1 month (session 3). As in session 1, session 3 was devoid of any trial-to-trial feedback.

To combine in our analysis the ‘enhance up’ and ‘enhance down’ groups, we now consider actions according to their congruency with the feedback. Specifically, 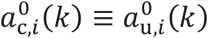for participants in the ‘enhance up’ group and 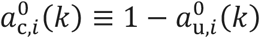 for participants in the ‘enhance down’ group. To quantify the global bias of a participant in a session, we averaged this quantity over the trials in that session. Because it took several trials for the bias to develop (Fig. 4A), we used the average of 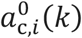 in the second half of the session 2, 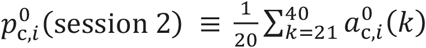 as a metric of the feedback-induced bias. For the bias session 3, we included all impossible trials: 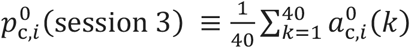.Omitting the index *i* for simplicity of notations, stability of preference would manifest as 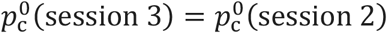 whereas 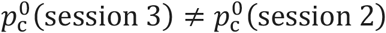 would imply that the preference of the participant has changed.

In our experiment, the delay between session 3 and session 2 was 1 day for one group of participants and 1 month for the other. Figure. 5A depicts 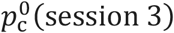 *vs*. 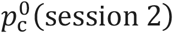 for each participant in the two groups. After 1 day, 25% of the participants (17/68) exhibited a significant change in 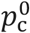 (closed circles in Fig. 5A, left). Of those, the number of participants who increased (*n*=7) and the numbers of participants that decreased their biases (*n*=10) were comparable (*p*=0.32, two-sided Binomial tests). This indicates that the feedback-induced bias was stable over one day. By contrast, in the one-month group, 29% of the participants (20/68) exhibited a significant change in their bias (closed circles), all of them with decreased it (*p*<0.00002, two-sided Binomial tests). These results indicate that the feedback-induced bias has significantly decayed within a month.

**Fig. 5.**
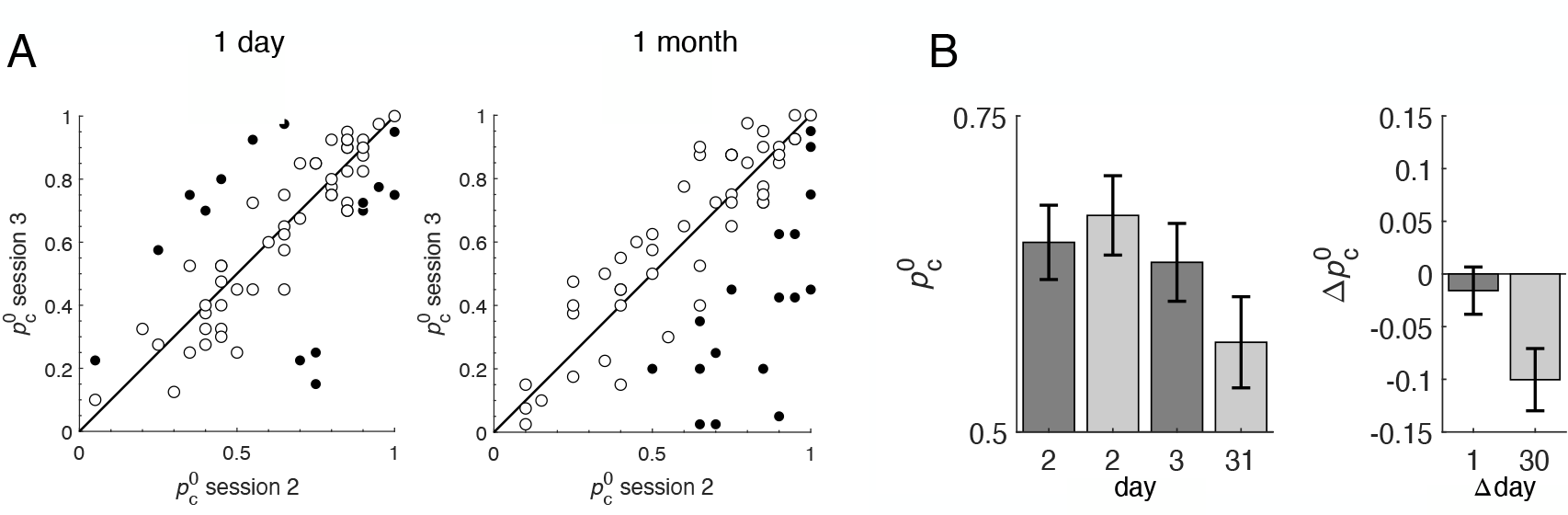
The feedback-induced bias is stable over one day but not over one month. **A**, the value of 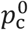 (the bias congruent with the feedback of session 2) in session 3 as a function of 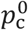 in session 2, in the one-day (left) or one-month (right) groups. Each dot corresponds to one participant. Filled circles: participants in which the change in 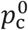 was statistically significant. Open circles participants with non-significant changes. In the 1-month group, 29% of the participants (20/68) exhibited a significant change in their bias with a decrease for all of them. **B**, Population analysis. The change in the population-average bias congruent with the feedback of session 2,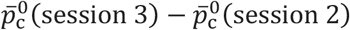. For the one-month interval group, 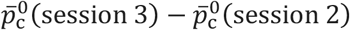 was significantly negative (*p* = 0.013, two-sided Wilcoxon signed-rank test).

To further quantify this decay, we considered the population-average of the bias. We define 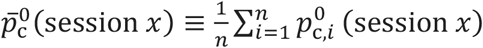, where *x* ∈ {2,3}. A decay of the feedback-induced bias would manifest as 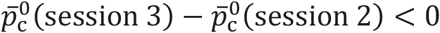 (Fig. 5B). For the 1-day group,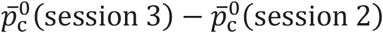 was not significantly different from 0 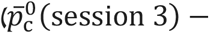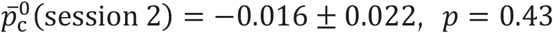, two-sided Wilcoxon signed-rank test). By contrast, for the 1-month interval group, 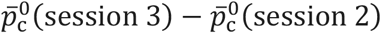 was significantly smaller than 0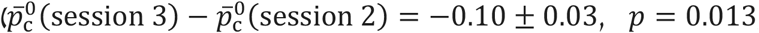, two-sided Wilcoxon signed-rank test). These results confirm that the feedback-induced bias which persists over 1 day, is unstable over 1 month.

To quantify the decay rate of the feedback induced bias, we considered the change in the global bias of the 1-month group from session 2 to session 3. Assuming that the decay is exponential, we can use the metric 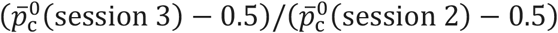 to quantify the decay time constant. We found 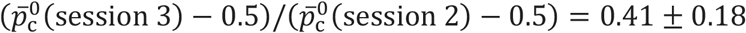 corresponding to a decay time constant of 1.1 months (bootstrap analysis shows that with 95% confidence, the decay time-constant is smaller than 2.6 months). Assuming this value, the feedback-induced bias is expected to diminish to 1% of its original value by 5 months. By contrast, the choice bias in the previous experiment remained stable for many more months (Fig. 2). Together, these results indicate that the feedback-induced bias is substantially less stable than the ICB observed in session 1.

To what value does the bias converge to at the *single participant* level? One possibility is that the feedback somehow resets the ICB. In that case, the correlation between the biases in sessions 1 and 3 will not increase with the time duration between sessions 2 and 3. Alternatively, the contribution of the feedback provided in session 2 will diminishes with time. In that case, the choice bias is predicted to revert with time to an ICB like the one in session 1.

Figure 6A plots the value of 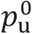 in session 3 *vs*. its value in session 1 for all the participants in the 1-day and 1-month groups (red and blue, ‘enhance up’ and ‘enhance down’ groups, respectively). Inspection shows that the dots are closer to the diagonal in the 1-month groups group suggesting that with time choice biases revert to their original value. To quantify this effect, we computed the Pearson correlation of 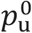 in sessions 1 and 3. Figure 5B) shows that this correlation is significant. It is smaller in the 1-day interval group compared with the 1-month interval group (Fig. 6B; *p* = 0.04, two-sided z-test for the difference in correlation after Fisher z-transformation). We thus conclude that at the single participant level the feedback-induced bias is transient, unlike the ICBs of session 1.

**Fig. 6.**
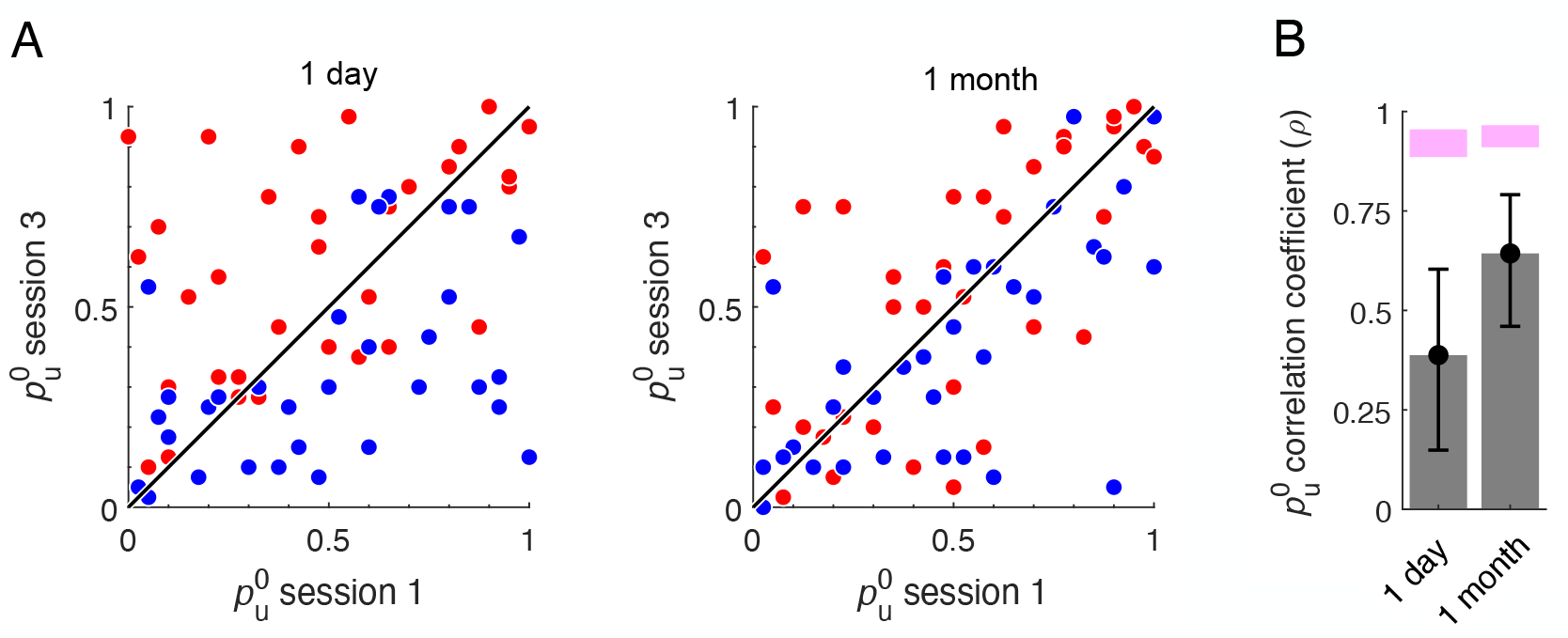
ICBs revert to their original values. **A**, value of 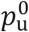 in session 3 as a function of 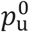 in session 1 for each participant in the 1-day (left) or 1-month (right) groups. Red dots: participants in the ‘enhance up’ group in session 2. Blue dots: and ‘enhance down’ participants in session 2. Line: the diagonal. **B**, Between-session correlation of 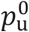 in sessions 1 and 3 for the two delay groups. Error bars: 95% confidence interval of the correlation, bootstrap. Magenta: 95% confidence interval of the correlation expected under the assumption of complete stability, bootstrap Bernoulli processes. The correlation between 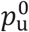 is significantly larger in the 1-month interval group than in the 1-day group (*p* = 0.04, two-sided z-test for the difference in correlation after Fisher z-transformation) indicating that as the feedback manipulation is ‘forgotten’, and the ICBs revert to their original values.

## DISCUSSION

We studied the long-term dynamics of human ICBs. We showed that in the absence of feedback, these ICBs are remarkably stable for many months. Moreover, we found that while they can be temporarily shaped using feedback, this effect is transient: the ICB exhibited before the feedback manipulation recovers within weeks. Together, our results challenge the hypothesis attributing ICBs to the idiosyncratic history of choices and reinforcers.

### The generality of the results

The focus of this paper was the vertical bisection trials. This is because previous studies have reported global biases in horizontal comparison tasks, which were attributed to pseudoneglect^24^. To test the generality of our results we interleaved the vertical bisection trials with trials of a different perceptual task. In the latter, participants were instructed to compare the sizes of two, horizontally displaced Gaussian-blurred circles (see Methods and Fig. S4A). Like the vertical-bisection task, 1/6 of the circle-comparison trials were impossible, and feedback about the correct answer followed the same schedule as in the vertical-bisection trials. The two types of trials were interleaved such that participants were presented with alternating blocks of three vertical bisection (B) trials and three horizontal comparison (C) trials such that the impossible trials (B^0^ and C^0^, respectively) were positioned as the first trial in a block (…BBBCCCB^0^BBC^0^CC…). This design was chosen as to minimize sequential effects in the impossible trials, which could confound our estimate of the ICBs in the impossible trials. This design also allowed us to test whether ICBs in the circle-comparison task are comparable to those of the bisection task and whether they follow the same dynamics following biased feedback. We replicated the main results of the paper in the circles’ task, namely, participants exhibited substantial ICBs, which were stable over many months. Feedback temporarily influenced these ICBs, but the induced effect was short-lived and decayed over time (Fig. S4-S9). This suggests that our main findings can be generalized to other perceptual comparison tasks.

### Relation to previous studies

Stability of ICBs has been demonstrated in humans over time intervals varying from days (*e*.*g*.^32^) to weeks (*e*.*g*.^11,13,33,34^). In ^11,33,34^, repeated measurements were used to quantify the stability of the bias, a procedure which may have a stabilizing effect on the bias^11^. Our study demonstrates that ICBs can be stable for much longer periods, at least two years, even without multiple measurements.

Of relevance to our work is a human study that used ambiguous stimuli to measure the stability over time of idiosyncratic biases in direction-of-motion perception^9^. Like our findings, biases in favor of a particular direction, measured at an interval of one year, were significantly correlated. However, exhibiting a substantial drift over time and measurements, these biases seemed to be less stable than those of our study.

Idiosyncratic side-preferences were also demonstrated in several species, including mice^16,17^, rats^18–20^, zebrafish^35^. In drosophila^15^, non-heritable idiosyncratic choice bias is stable for at least one month, a substantial fraction of their 2-3 months lifespan.

Our finding that participants are sensitive to feedback is in line with studies in a vernier and 2-tone discrimination tasks^25,36^. They reported that providing incorrect feedback for a subset of high difficulty or impossible trials rapidly biases choices congruently with its asymmetry. In the vernier task, however, the effect of the feedback decays within the session (timescale of minutes), once the feedback is turned off^25^, and is not transferred overnight^26^. Along the same lines, providing correct trial-by-trial feedback to participants that exhibit ICB can reduce it within the experimental session^37^. Beyond perceptual tasks, choice preference induced by feedback is a form of operant learning^38,39^, and can have long-lasting effects.

### Limitations of the study

Our conclusion that ICB is not the result of idiosyncratic feedback-induced learning is based on the discrepancy between the relatively transient nature of the feedback-induced bias and the stability of the ICB. However, we cannot exclude the possibility that this discrepancy reflects the specific conditions in which we induced the bias. For example, it is possible that feedback-induced bias, if applied at an earlier age, would have led to a more sustained bias. It is also possible that providing biased feedback over more trials and / or for more sessions would have resulted in a longer-lasting effect^25–31^. Moreover, it is possible that the dynamics of this decay are characterized by multiple timescales. In this case, the fact that feedback-induced bias *decays* within weeks does not imply that it would almost *fully* disappear if we waited months. Perhaps some of the timescales that characterize the dynamics of the decay are sufficiently long to account for ICB.

### What underlies ICBs and their stability

Previous studies have shown that choices are strongly affected by the history of stimuli that the participant has been exposed to, a phenomenon that can be explained in the Bayesian framework^40^. Therefore, a potential contributor to the ICB is the idiosyncratic distribution of stimuli that the participants were exposed to. However, this contribution is likely to be small. First, participants are primarily affected by the most recent stimuli^41^ whereas in our experiments, a distribution of ICBs is observed even though within the experiment all participants were exposed to the same sequence of stimuli. Second, the influence of perturbing stimuli on bias has been shown to decay within tens of seconds^9^.

Another possible contributor are stable idiosyncratic asymmetries. For example, asymmetries in the anatomy of the eyes could give rise to asymmetries in the perception of the bar segments, which would manifest in a stable perceptual bias. While such idiosyncratic anatomical asymmetries are likely to exist, several lines of evidence suggest that this contribution is not dominant. First, we have previously reported a comparable distribution of ICBs in a sensory-motor task that does not entail any sensory ambiguity^14^. Second, in drosophila, left-right ICB and idiosyncratic object orientation control have been shown to be uncorrelated with anatomical asymmetries (e.g., between the legs length)^15,42,43^. Finally, if the idiosyncratic asymmetry is weak relative to the width of the psychometric curve, its contribution to the bias would be negligible. Alternatively, if it is relatively strong, participants would always report ‘up’ or always report ‘down’ when the two segments are of the same length. Only a small range of idiosyncratic asymmetries is consistent with the experimentally observed wide distribution of ICBs.

In a previous study^14^, we showed that under general conditions, a wide distribution of ICBs emerges naturally from the dynamics of competition between neuronal networks that are *statistically identical*, that is networks whose parameters are drawn from the same distribution. The intuition behind this result is that in such networks the microscopic structure of the connectivity varies across networks because they are only *statistically* identical. The main result there was that surprisingly, such heterogeneity is predicted to substantially bias choices even in (infinitely) large decision-making networks. If the ICBs are the result of microscopic heterogeneities in the synaptic connections of decision-making networks, the long-term dynamics of the former can be related to one of the latter. We do not know much about the stability of the synaptic connectivity in decision-making networks, certainly not in humans. However, the question of stability of connectivity has been previously addressed, primarily by spine imaging of neurons in the mouse cortex. Several studies have shown that in some regions the connectivity is highly volatile^44^. For example, in the mouse auditory cortex, most dendritic spines on pyramidal neurons, a proxy of the excitatory synapses that reside on them, are replaced within three weeks^45^. Of the remaining spines, most change their size by at least a factor of two within that period^46^. These results indicate that the fine structure of connectivity is highly dynamic^44^, which according to our mechanism predicts a substantial volatility of ICBs in a timescale of weeks. On the other hand, in the barrel cortex, a fraction of spines is stably maintained for months in support of a stable connectome^47^ and stable ICBs. The stability of the ICBs in our experiments predicts stability of the underlying connectivity.

## Methods

The experiments were approved by the Hebrew University Committee for the Use of Human Subjects in Research and all participants provided informed consent. Recruitment was based on the online labor market Amazon Mechanical Turk^38^. Each of the studies was described as a longitudinal academic survey of visual acuity.

### Study 1: Stability of ICB in absence of feedback

Human participants were invited to take part in a two-session experiment. Upon the completion of the first session (session 1), they received a link indicating the time-window of the second session. Of a total of 245 participants, 62 who failed to start the second session (session 2) within their designated time window were excluded. This resulted in a final sample size of 183 participants, as described in Table 1. In addition, 88 of those who completed session 1 and session 2 in 8 days or less were invited to participate in a third session (session 3), 22 months later; 29 of them completed this session (see: Table 1, bottom row). None of the participants that started session 2 on time were excluded from the analysis.

**Table 1.** Study 1 – demography of participants included in the analysis.

| Time between sessions |  | *n* | # females | # sinistral | Age on session 1 (years) |  |
| --- | --- | --- | --- | --- | --- | --- |
| Mean | Range |  |  |  | Mean | Range |
| 1.7 hours | 0.3-2.6 hours | 29 | 19 | 3 | 39.9 | 24-67 |
| 26.3 hours | 23.1-42.9 hours | 32 | 24 | 4 | 41.0 | 28-69 |
| 6.7 days | 6.0-7.9 days | 27 | 13 | 3 | 40.4 | 24-66 |
| 28 days | 27-31.9 days | 33 | 18 | 1 | 38.8 | 22-61 |
| 88.5 days | 85.9-97.1 days | 30 | 18 | 3 | 39.6 | 25-65 |
| 240.1 days | 232.2-247.2 days | 32 | 15 | 2 | 40.9 | 24-69 |
| 665.7 days | 656.7-672.1 days | 29 | 22 | 2 | 42.3 | 28-66 |

A base monetary compensation was given to all participants. separately in each session. To encourage good performance, an additional bonus fee was given for every correct response and another bonus was guaranteed to 10% of participants with highest scores in each session. All participants were in the United States of America and all reported normal or corrected to normal vision.

#### Experimental procedure

All sessions followed the same procedure. In each trial, participants were instructed to make a perceptual discrimination decision. There were two types of perceptual tasks, a vertical bisection task, analyzed in the Results section and a horizontal, size-comparison task, which is mentioned in the Discussion section and the Supplementary Information. Participants were instructed to make their decision as quickly and as accurately as possible.

In a bisection trial, a 200 pixel-long vertical white line, transected by a horizontal 20 pixel-long white line was presented on a black screen and participants were instructed to indicate which segment out of two is longer (Fig. 1A, inset). In a size-comparison trial, two white Gaussian blur circles with an average radius of 75 pixels were presented on a black screen and participants were instructed to indicate which circle out of two is bigger (Fig. S1A, inset). In both cases, participants were instructed to press the spacebar key once coming to a decision. Upon the spacebar press, the stimulus was replaced by a decision screen composed of two arrows buttons, appearing on opposite sides of the screen, and a middle 4-squares submit button. In the bisection task, the two arrows were either ‘up’ or ‘down’, whereas in the size-comparison task, they were either ‘left’ or ‘right’. The participants indicated their decision by moving the initially centered cursor to one of the arrow buttons, pressing it, and finalizing their decision by pressing the ‘submit’ button. Upon the ‘submit’ button press, the decision screen was replaced by a black screen and the next trial began after 500 milliseconds.

The stimuli were limited to a 400-pixel x 400-pixel square at the center of the screen. The horizontal location of all vertical bisection lines and the vertical location of the center of all circles were centered. Window resolution was verified for each participant individually, to make sure that it did not exceed the centric box in which all stimuli were presented.

No feedback was given regarding the correct response. The participants were, however, informed about the accumulated bonus fee every 30 trials.

Each session consisted of 480 trials, 240 vertical bisection (B) and 240 horizontal size-comparison (C) trials. Trials in each session were ordered in 160 alternating blocks of 3 bisection and 3 size-comparison trials (…CCCBBBCCCBBB…). Unknown to the participants, there were 40 impossible bisection and 40 impossible size-comparison trials (⅙ of the trials) in each session. To minimize sequential effects in the impossible trials, each impossible bisection trial was preceded by three size-comparison trials. Similarly, each impossible size-comparison trial was preceded by three bisection trials. For the possible bisection trials, the deviation from the veridical midpoint was uniformly distributed between 2, 4, 6, 8 and 10 pixels. The quantity *L*^U^ and *L*^*D*^ denoting the lengths of the ‘up’ and ‘down’ segments of the vertical line, this corresponds to 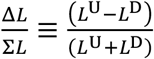 between 0.02 and 0.1, with an equal number of offsets in each direction. For the possible size-comparison trials, the difference in radii between the two circles was uniformly distributed between 4, 8, 12, 16 and 20 pixels. Defining 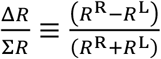with *R*^R^ and *R*^L^ the radii of the right and left circles this corresponds to 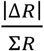 between 0.0267 and 0.1333, with an equal number of deviations in each direction. To minimize external causes of heterogeneity, the order of the trials was identical for all participants.

To verify that the participants understood the instructions, they were required to successfully complete a bisection practice session and a size-comparison practice session prior to the experiment. This session consisted of blocks of 4 easy trials with feedback and balanced polarity of Δ*L* or Δ*R*. In each session, the main experiment started after the participant completed one bisection and one size-comparison block successfully. Responses in this practice session were not included in the analysis.

Mean performance in the possible bisection trials was 92.1% ± 5.3% (standard deviation), range 62% − 99.5% in the first session and 92.6% ± 5.4% (standard deviation), range 66% − 99.5% in the second session. Mean performance in the possible size-comparison trials was 98.6% ± 1.8% (standard deviation), range 89.5% − 100% in the first session and 98.3% ± 3.6% (standard deviation), range 58% − 100% in the second session.

### Study 2: Feedback-induced bias and its stability

This study was like study 1 in terms of tasks, stimuli and instructions, with one major exception: it consisted of intermediate feedback third session (session 3). Participants were told in advance that the strict time window for starting session 2 is 24-54 hours after completion of session 1. In addition, participants were told in advance that they will be informed about the time window for starting session 3 only during session 2. The time window could either be 24-72 hours or 28-35 days after completion of session 2. This allowed us to match participants into groups based on their ICBs in session 1 (see below) to create groups that are as possible similar in their initial biases. Overall, data in session 1 were collected from 148 participants. We excluded from the analysis those participants who failed to start the session 2 and session 3 within their designated time windows (see below, 12 participants, 8.1% dropout). This resulted in a final sample of 136 participants.

After completing session 1 and before starting session 2, all participants were divided into groups that later defined both their feedback manipulations in session 2 and the time window between the session 2 and session 3. The feedback manipulations in session 2 were either ‘enhance up’ or ‘enhance down’ for the bisection task and either ‘enhance right’ or ‘enhance left’ for the size-comparison task. Hence, each participant was associated with one bisection manipulation group and with one size-comparison manipulation. Further, half of the participants waited 1-3 days between session 2 and session 3 whereas the other half waited 28-34 days. Thus, there were 8 groups of participants. Because our studies primarily focused on the bisection task, participants were first divided into the bisection manipulation and delay time groups according to their 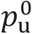,as measured from the first session, such that each of the four groups (‘enhance up’, ‘enhance down’ × 1-day, 1-month) had as similar distribution of initial 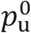 biases as possible. Specifically, we first sorted in ascending according to 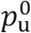 of all the 148 participants in session 1. These were further divided into 37 quartets, each quartet consisting of 4 participants with consecutive 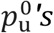. Following, the participants in each quartet were randomly divided between the 4 groups of bisection ‘enhance’ polarities and delay times (1 participant per group).

This routine was repeated until all participants were assigned to a bisection ‘enhance’ and a time delay group. Only then were the participants in each of the 4 groups above further divided into size-comparison manipulation groups, based on matching their 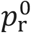, defined as the fraction of responses that the circle to the right was larger, in session 1. Table 2 provides a summary for each group.

**Table 2.** Study 2 – Participants included in the analysis.

| Table 2. Study 2 – Participants included in the analysis. |  |  |  |  |  |  |  |  |
| --- | --- | --- | --- | --- | --- | --- | --- | --- |
| Feedback manipulation in session 2 |  | Time between sessions 1 and 2 |  | n | # females | # sinistral | Age on first session (years) |  |
|  |  | Mean (days) | Range (days) |  |  |  | Mean | Range |
| ‘Enhance up’ | ‘Enhance right’ | 1.2 | 1-1.9 | 14 | 6 | 2 | 40.1 | 25-70 |
|  |  | 29.3 | 28-33.9 | 18 | 6 | 3 | 36.5 | 19-54 |
|  | ‘Enhance left’ | 1.1 | 1-1.8 | 20 | 8 | 3 | 40.8 | 26-60 |
|  |  | 28.5 | 28-29.9 | 17 | 7 | 2 | 42.2 | 23-74 |
| ‘Enhance down’ | ‘Enhance right’ | 1.2 | 1-2.1 | 18 | 6 | 5 | 38.3 | 22-68 |
|  |  | 28.5 | 28-31.3 | 17 | 7 | 6 | 44.4 | 27-72 |
|  | ‘Enhance left’ | 1.2 | 1-2.9 | 16 | 7 | 2 | 41.5 | 25-60 |
|  |  | 29.5 | 28.2-32.9 | 16 | 8 | 0 | 39.7 | 28-55 |

As in study 1, participants received a link with a countdown for the time to start the next session and were informed about the time window. Once the countdown was over, participants were automatically invited to take part in the next session. Base monetary compensation and bonuses were given to all applied participants in each session (separately). In addition, to encourage participants to take part in all three sessions, an additional bonus was given to those who completed session 3. As in study 1, all participants were in the United States of America. They reported normal or corrected to normal vision.

#### Experimental procedure

The sessions in this study consisted of the bisection and size-comparison tasks described above for study 1, with the same number of impossible and possible bisection and size-comparison with one exception. For the possible size-comparison trials, the difference in radii between the two circles was smaller than in study 1 (trials were more difficult): uniformly distributed between 2, 4, 6, 8 and 10 pixels (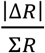 between 0.0133 and 0.0667).

Session 1 and session 3 followed the same procedure as in study 1. Session 2 differed in that after each of the 480 trials participants received feedback about their response, a smiley or a sad emoji indicating whether their choice was correct or incorrect. The feedback was accurate in all possible trials (Δ*L* ≠ 0 and Δ*R* ≠ 0, 5/6 of the trials). By contrast, in the impossible trials, different feedback schedules were used. For participants in the ‘enhance up’ group, the ‘up’ response was considered ‘correct’ in 95% of the impossible trials, whereas for the ‘enhance down’ group, the ‘down’ response was considered ‘correct’ in 95% of the impossible trials. For participants in the ‘enhance left’ group, the ‘left’ response was considered ‘correct’ in 95% of the impossible trials, whereas for the ‘enhance right’ group, the ‘right’ response was considered ‘correct’ in 95% of the impossible trials (see Table 2).

## Data and code availability

Datasets are available on the Zenodo archive (https://doi.org/10.5281/zenodo.13388598).

MATLAB code used for all analyses is available on the Zenodo archive (https://doi.org/10.5281/zenodo.13388598) and in the stabilityFeedback GitHub repository (https://github.com/Lior-Lebovich/stabilityFeedback).

## Acknowledgements

This work was supported by the German Research Foundation (DFG CRC 1080) and the Gatsby charitable foundation (Y.L) and in part by ANR-17-NEUR-005 (DH). Y.L. is the incumbent of the David and Inez Myers Chair in Neural Computation. Work performed in the framework of the France-Israel Center for Neural Computation (CNRS/Hebrew University of Jerusalem). L.L. thanks Nelly Lebovich for insightful discussion.

## Author contributions

L.L., L.K., D.H. and Y.L. conceived and planned the experiments; L.L. and L.K. programmed and ran the experiments; L.L. and Y.L. analyzed the data; L.L., D.H. and Y.L. wrote the manuscript.

## Supplementary Information

**Fig S1.**
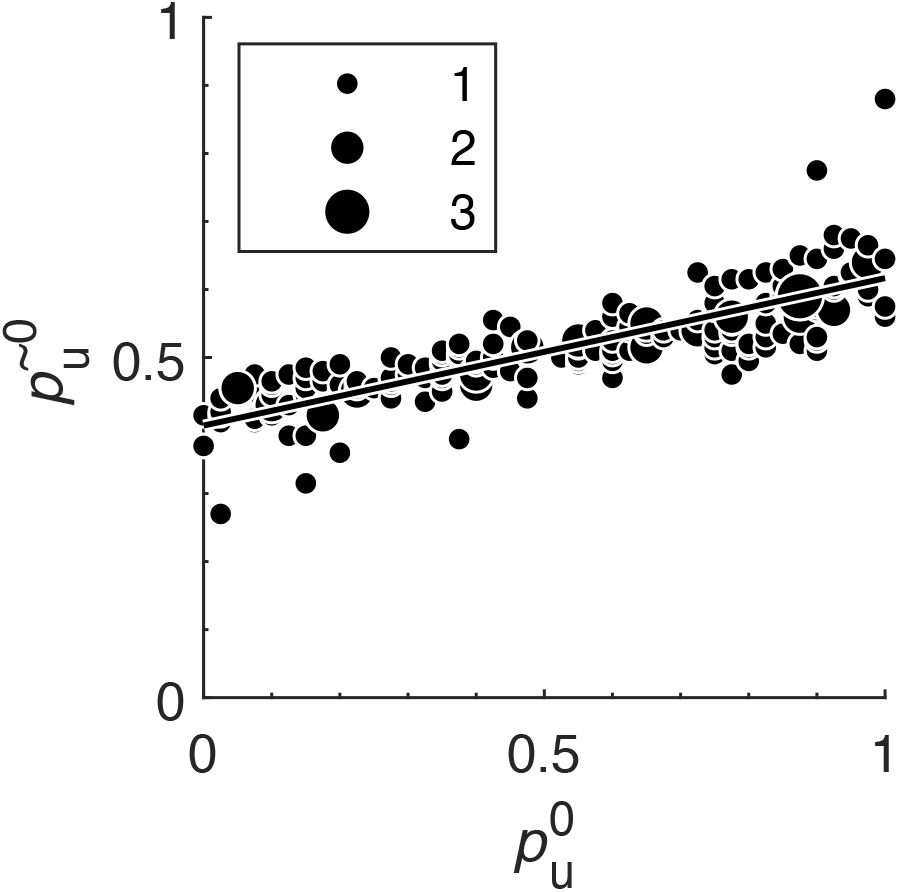
ICBs in the impossible and possible trials are correlated in the vertical bisection task. For each of the 183 participants, the fraction of ‘up’ responses was computed separately for the possible and impossible trials. Each dot denotes the value of *p*_u_ in the possible trials 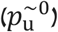 *vs*. its value in the impossible trials 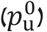. Small dots: single participants. Radii of larger dots: the number of participants sharing the same values of both possible and impossible *p*_u_ (see inset). Black line: best fit orthogonal regression (slope = 0.22). Despite the overall high performance in the possible trials (92.1% ± 5.31%), participants exhibited substantial ICBs in these trials, which were strongly correlated with the ICBs in the impossible trials (*ρ* = 0.82, p<0.001, *n* = 183 participants).

**Fig. S2.**
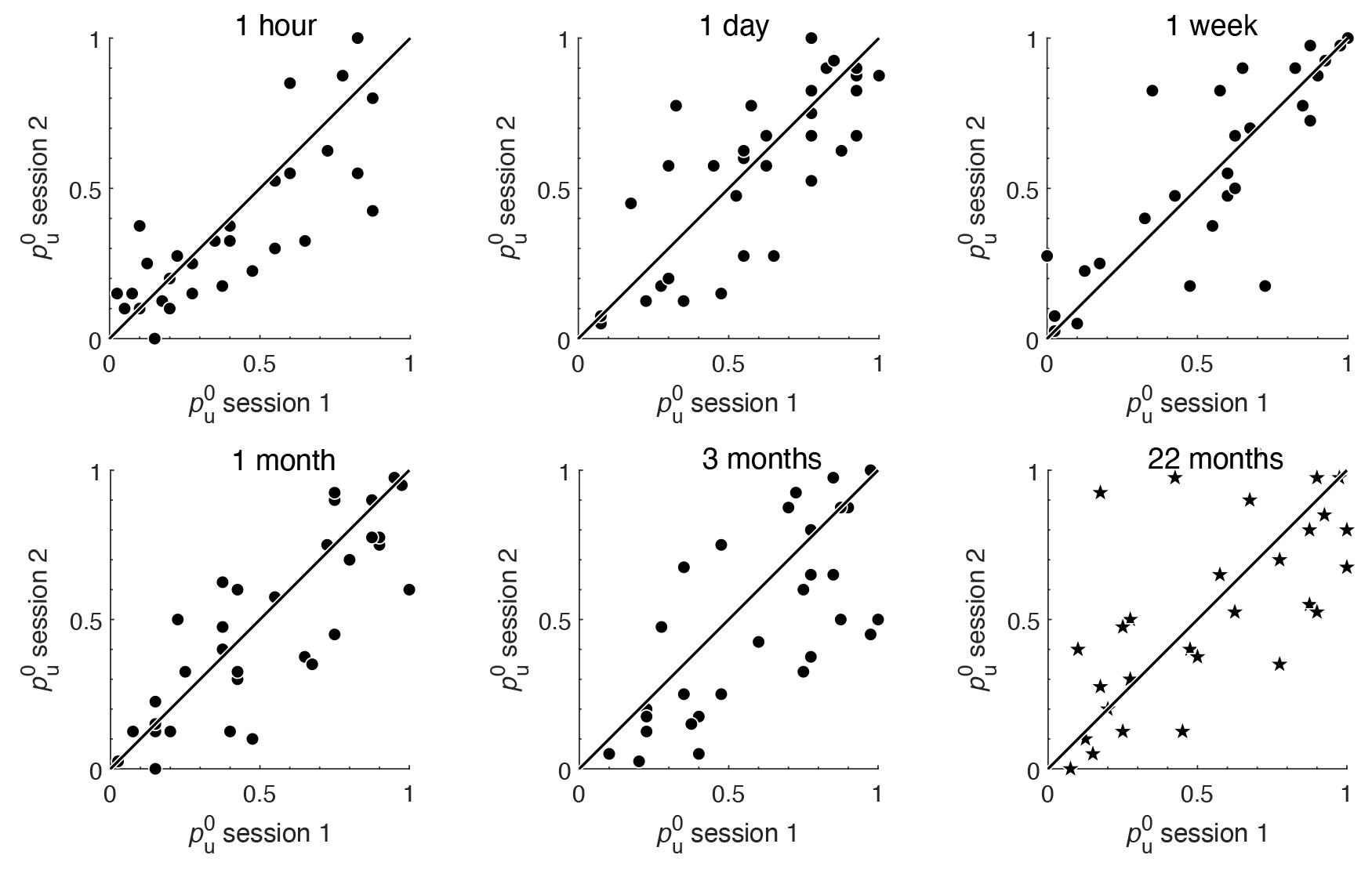
Stability of ICB by time-interval group in the vertical bisection task. Same as in Fig. 2B for participants who waited **A**, 1 hour, **B**, 1 day, **C**, 1 week, **D**, 1 month and **E**, 3 months between sessions 1 and session 2. **F**, Same analysis for the 22 month-interval between session 2 and session 3.

**Fig. S3.**
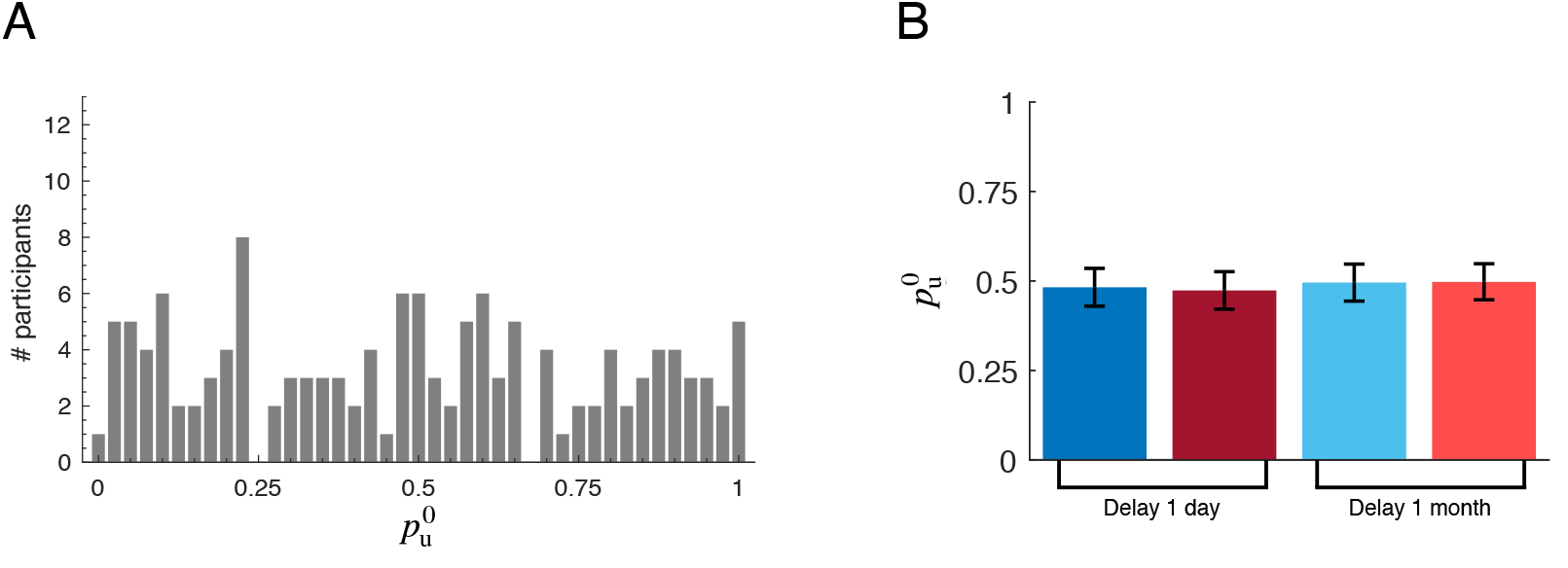
ICBs in the first session of the vertical bisection task. **A**, The distribution of ICBs in the first session, 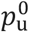, across all participants (*n* = 136). Overall, 64% of the participants exhibited a significant choice bias (29% significant 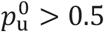 and 35% significant 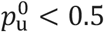; *p* < 0.05, two-sided Binomial tests, not corrected for multiple comparisons). As in Fig. 1B, the variance of the distribution is significantly larger than expected by chance (*p* < 0.001, two-sided bootstrap test, Bernoulli process; *n* = 136 participants, 40 trials per participant). While individual participants exhibited substantial biases, we could not detect any global bias at the population level: the average 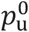 across all participants was 0.49, not significantly different from chance (95% CI 0.44-0.54, bootstrap). **B**, Matched groups based on ICB in the first session. The sample was divided into 4 groups that will later differ in the feedback manipulation in session 2 (Red and Blue color: ‘enhance up’ and ‘enhance down’ groups, respectively) and by the expected delay between the session 2 and 3 (abscissa). Bars: each group’s average 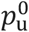 in all 40 impossible trials of session 1. Error bars: standard error of the mean. The 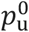 in all groups were comparable (p=0.99, Kruskal–Wallis one-way analysis of variance on ranks) and at the population level, no group exhibited any bias (delay and manipulation groups, from left to right: *p* = 0.60, *p* = 0.76, *p* = 0.93, and *p* = 0.97, two-sided Wilcoxon signed-rank test for the group’s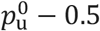, not corrected for multiple comparisons).

**Fig. S4.**
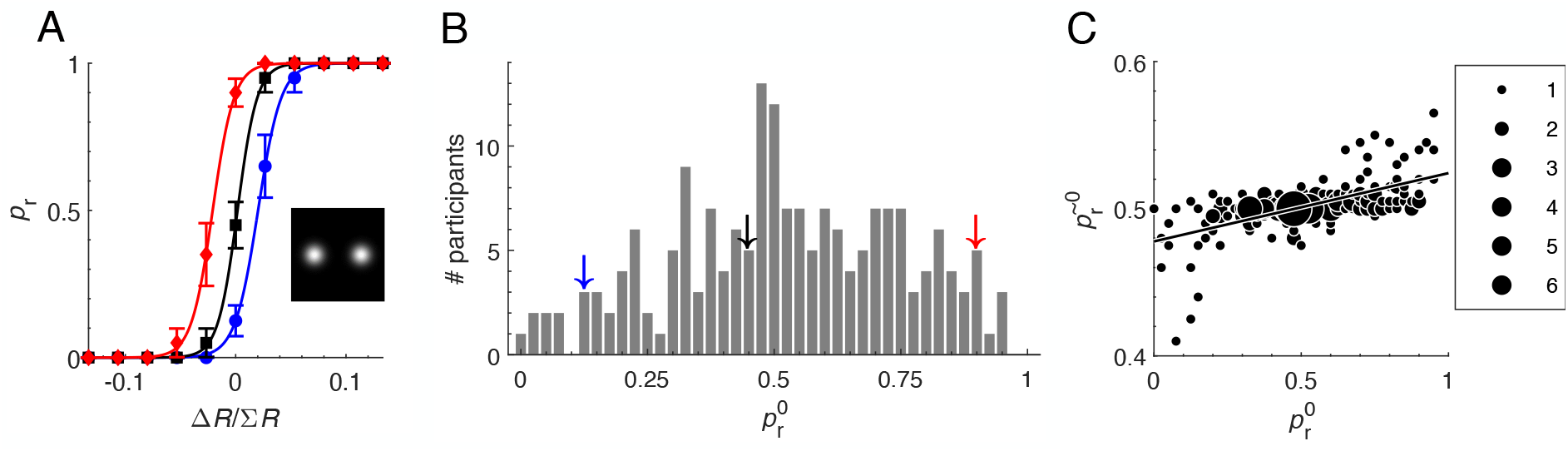
The horizontal comparison task. While the horizontal-circle size comparison task (A, inset) was mainly intended to reduce sequential effects, we repeated the analyses presented in the Result section of the paper for the horizontal task. **A**, Psychometric curves of three participants. The observed fraction of responding ‘right’,, is plotted *vs*. the sensory offset 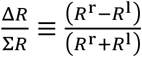, *R*^r^ and *R*^l^ denote where the radii of the right and left circles. Error bars: the standard error of the mean (SEM). Curves are best-fit logistic functions. Notably, the 3 example participants are different from the ones in Fig. 1A. **B**, The distribution across the participants (*n* = 183) of 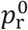, the value of *p* in impossible trials in (Δ*R* = 0). Arrows: the value of 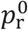 for the three participants in panel A, color coded. **C**, ICBs in the impossible and possible trials are correlated. For each of the 183 participants, the fraction of ‘right’ responses was computed separately for the possible and impossible trials. Dots: the value of ICB in the possible trials as a function of its value in the impossible trials. Small dots: single participants. Radii of larger dots denote the number of participants sharing the same values of both possible and impossible ICBs (see inset). Black line: best fit orthogonal regression (slope = 0.05). Despite the overall high performance in the possible trials (98.6% ± 1.8), participants exhibited ICBs in the possible trials that were correlated with the ICBs in the impossible trials (*ρ* = 0.61, *p* < 0.001, *n* = 183 participants).

**Fig. S5.**
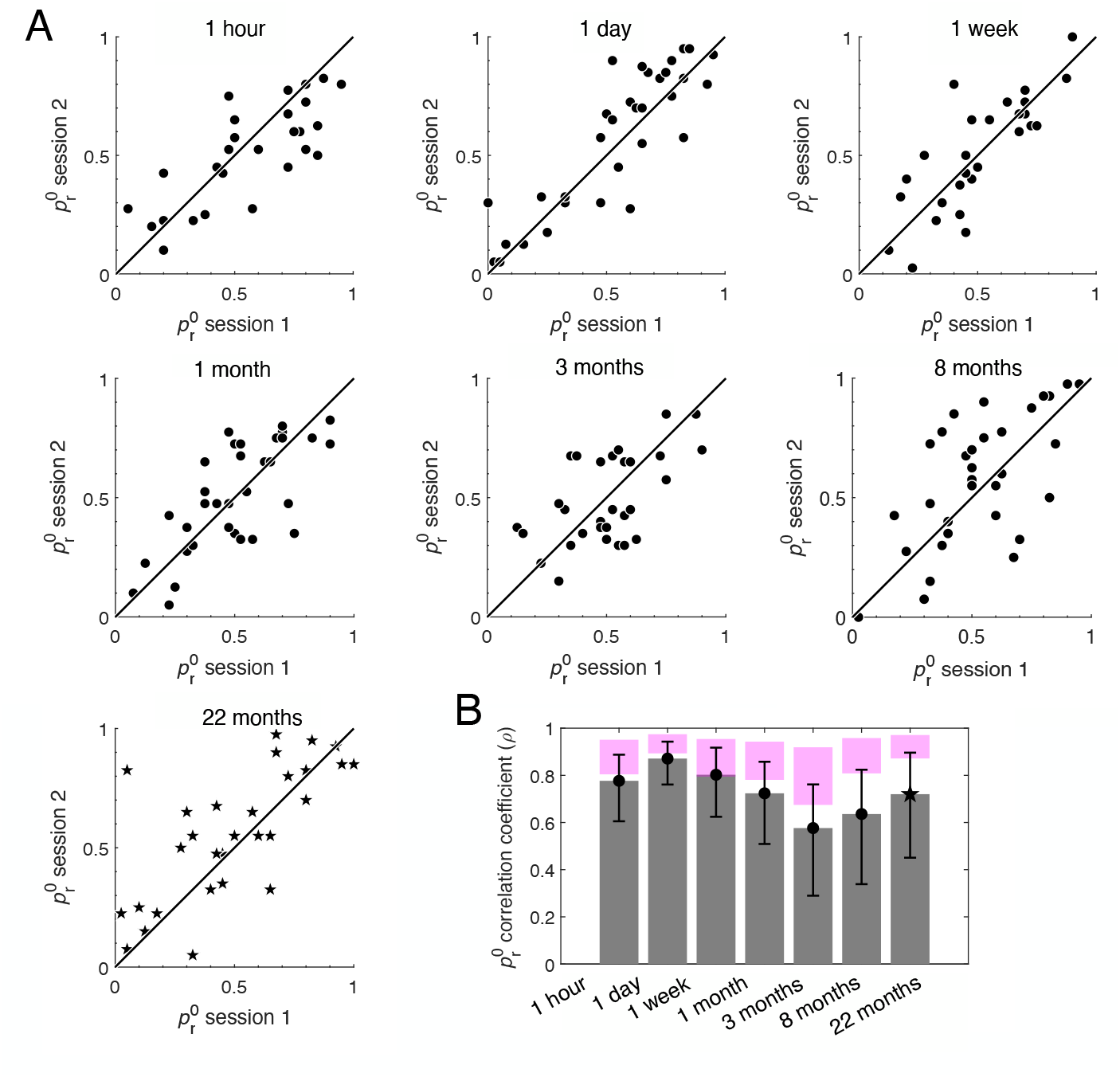
Horizontal ICB is stable over time. **A**, Same as in Fig. 2B and Fig. S2 for the horizontal comparison task. **B**, Same as in Fig. 2C for the horizontal comparison task.

**Fig. S6.**
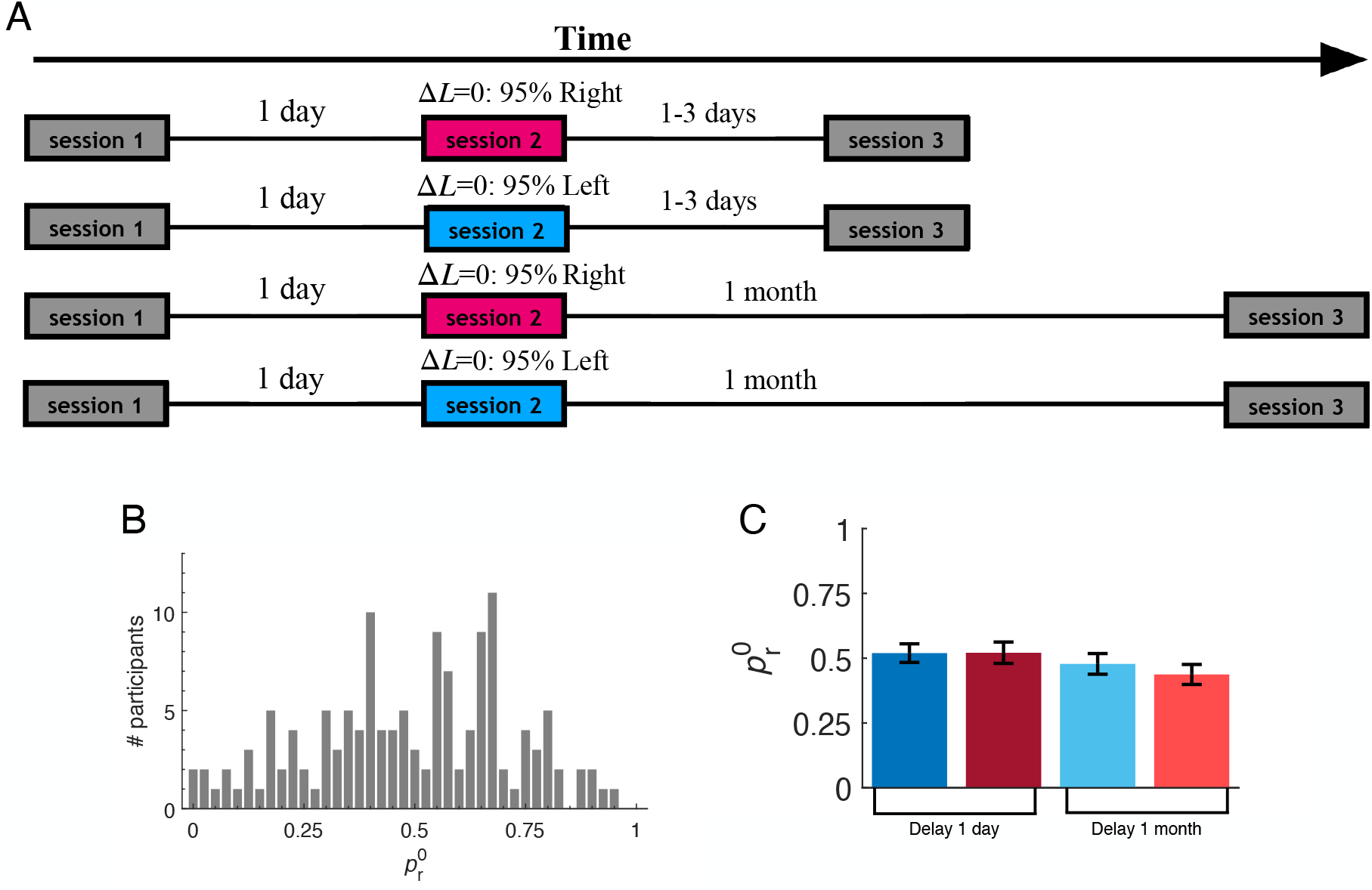
A, Design of the feedback-induced bias for the horizontal comparison task. **A**, Schematics of experimental design. All participants (*n* = 136) took part in three experimental sessions. Gray: there was no trial-by-trial feedback about the correct response in session 1 and session 3. In session 2, choices in all trials were followed by feedback. Feedback in the possible trials (200 trials) was accurate. Impossible trials (40 trials) were followed by asymmetrical feedback about the ‘correct’ response. Red: ‘enhance right’ group of participants, receiving feedback that the ‘correct’ response is ‘right’ in 38 impossible trials and ‘left’ in 2 impossible trials. Blue: ‘enhance left’ group of participants, receiving feedback that the ‘correct’ response is ‘right’ in 2 impossible trials and ‘left’ in 38 impossible trials. All participants took part in the second session 1-2.2 days (range; median: 1.02 day) following their completion of the first session. Half of the participants took part in session 3, 1-2.9 days (range; median: 1.03 day) following completion of session 2 (Top). The other half participated in session 3, 28-33.9 days (range; median: 28.4 days) following completion of session 2 (Bottom). Note that the sample was first divided, based on vertical ICB, into 4 groups which will later differ in the vertical-bisection feedback manipulation and the expected delay between session 2 and session 3 (see: Methods). Only then, was each of the 4 groups further divided based on their initial ICBs in the horizontal-comparison task into 2 groups that will differ in the feedback manipulation in session 2. **B**, The distribution of 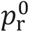 across all participants (*n* = 136). **C**, Matched groups based on horizontal ICB in session 1. Red and Blue color: ‘enhance right’ and ‘enhance left’ groups, respectively. Abscissa: expected delay between the session 2 and session 3. Bars: average 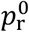 in all 40 impossible trials of the first session for each group. Error bars: standard error of the mean. The value of 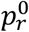 in all groups were comparable (p=0.38, Kruskal–Wallis one-way analysis of variance on ranks) and at the population level, no group exhibited any bias (delay and manipulation groups, from left to right: *p* = 0.51, p=0.48, *p* = 0.16, and *p* = 0.77, two-sided Wilcoxon signed-rank test for the group’s 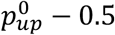 median, not corrected for multiple comparisons).

**Fig. S7.**
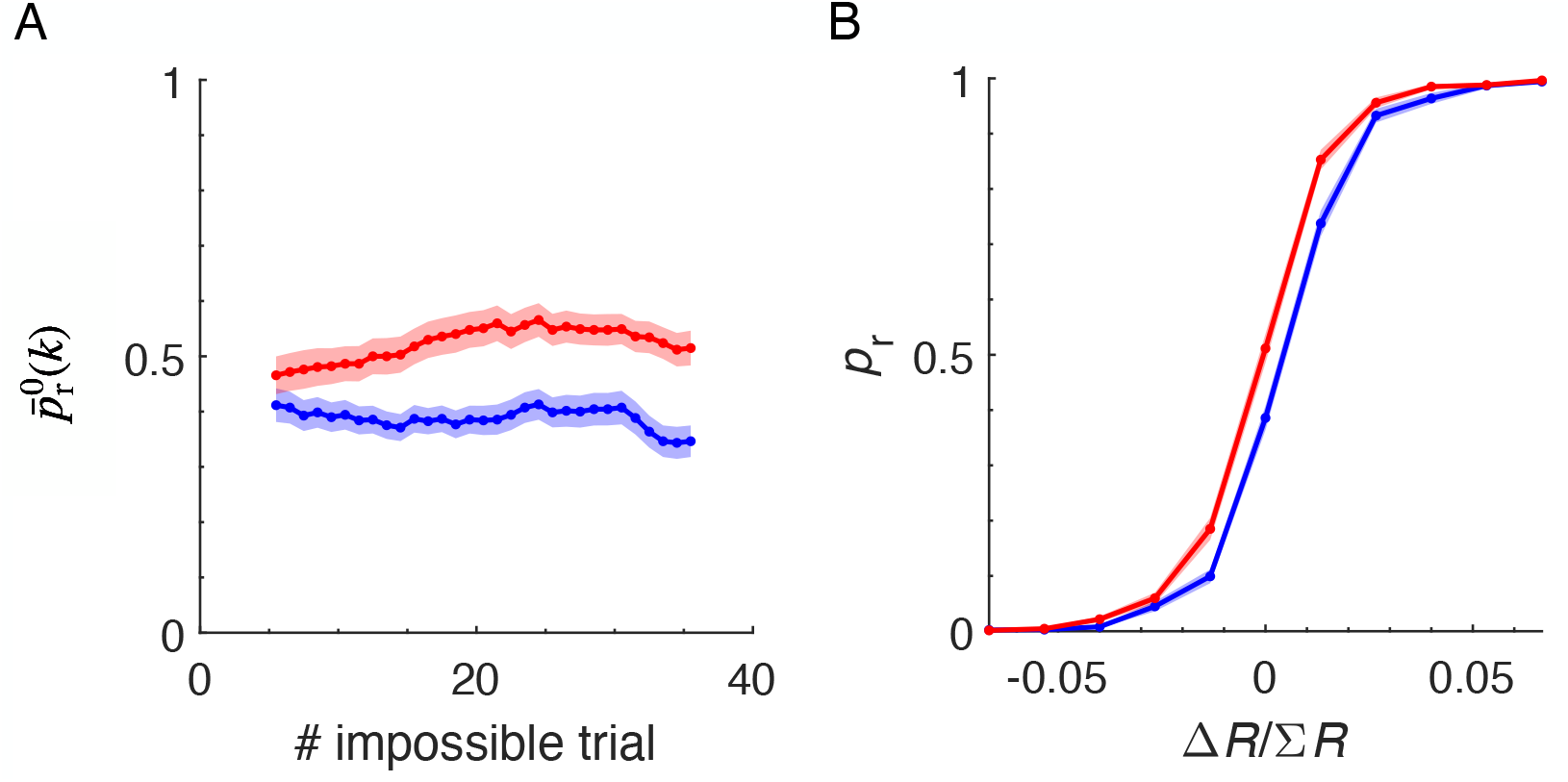
Effect of the feedback in the horizontal trials of session 2. **A**, the feedback effect in impossible trials. Group average of 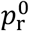, in a running window of 10 impossible trials (dot *k* denotes the group average 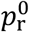 in impossible trials *k* − *k* + 9), similar to Fig. 4A. Red, ‘enhance right’ participants. Blue, ‘enhance left’ participants. Shaded area: SEM. **B**, the effect of the feedback in possible trials. Group average psychometric curve (line) and SEM (shaded area) of *p* as a function of 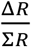, the relative difference between the radii of the two circles, like Fig. 4B. Color coded as in **A**.

**Fig. S8.**
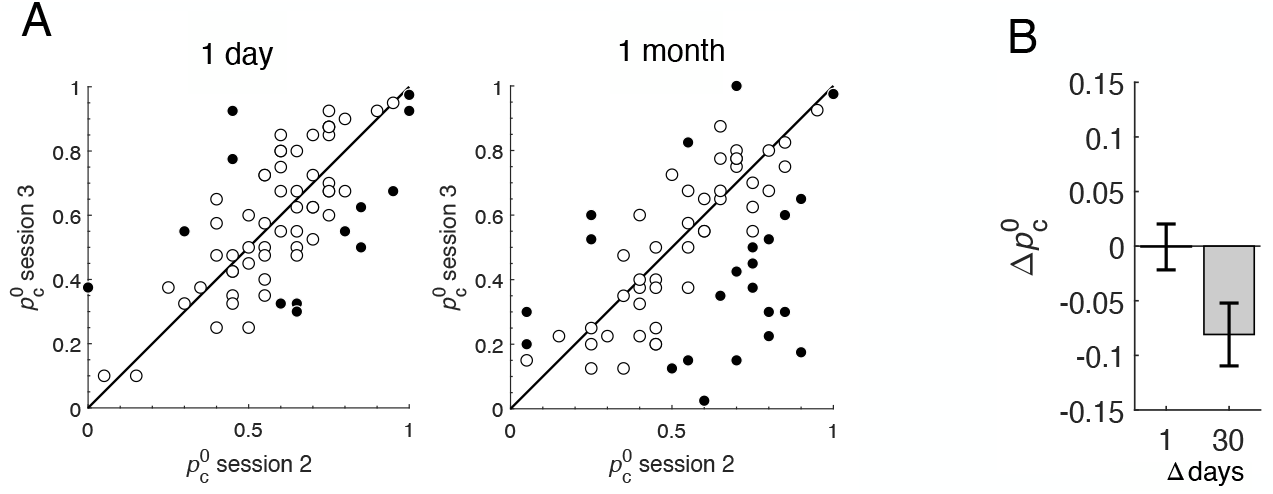
The decay of the effect of the feedback in the horizontal task. **A**, each dot depicts a single participant who waited 1-day (left) or 1-month (right) between session 2 and session 3. 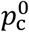 is defined by the manipulation in session 2; namely: 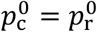 for participants in the ‘enhance right’ groups and 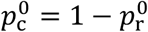 in the ‘enhance left’ groups. Abscissa: fraction of congruent responses in the second half of session 2,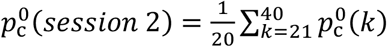. Left: 1-day interval. Right: 1-month interval. Filled circles: participants in which the change in 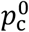 was statistically significant. Open circles: participants for which it was not significant. B, population analysis. The change in the population-average bias congruent with the feedback of session 2, 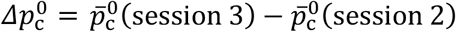.Error bars are SEM. While the average 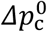 (1 day) was not significantly different from 0 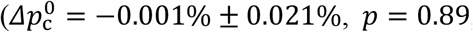, two-sided Wilcoxon signed-rank test), the average 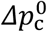 (1 month) was significantly negative 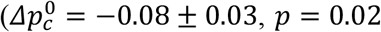, two-sided Wilcoxon signed-rank test), indicating that the feedback-induced effect is substantially less stable than the horizontal ICBs measured in Fig. S5.

**Fig. S9.**
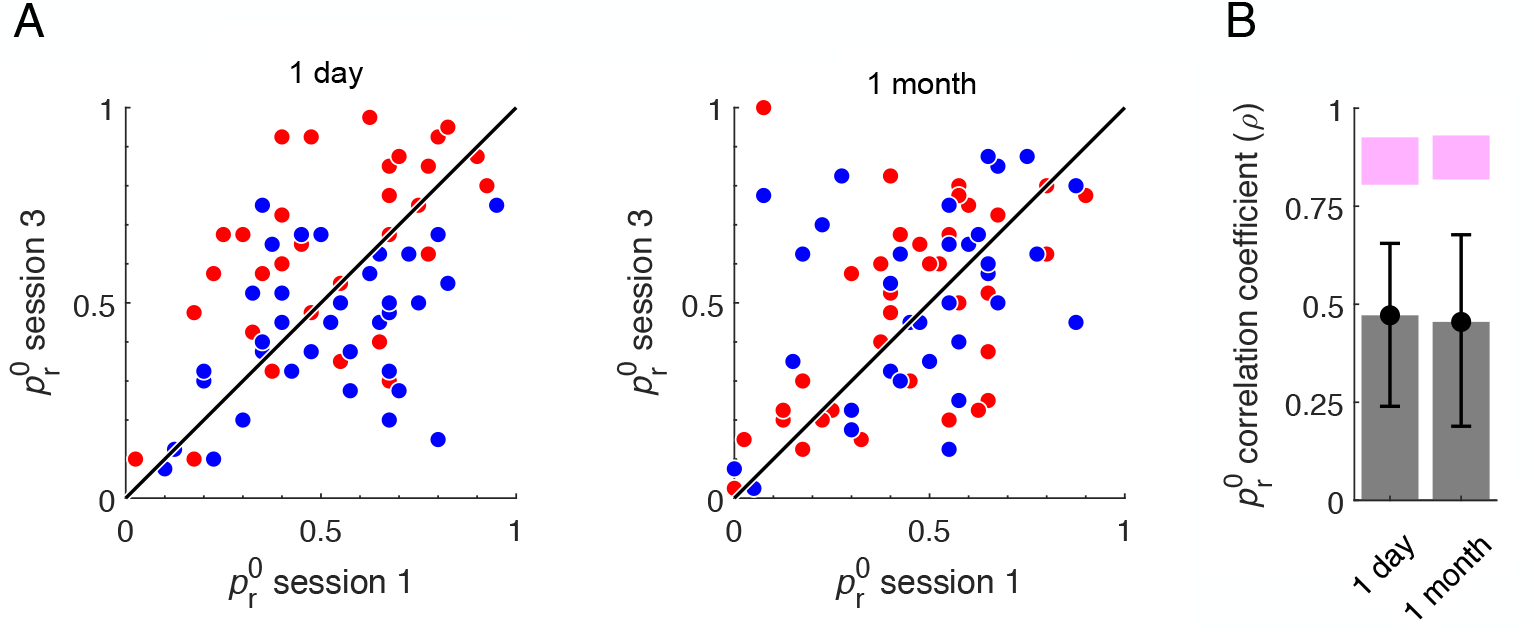
Temporal dynamics of the feedback effect on individual horizontal ICBs. **A**, Dots: 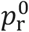 in session 3 as a function of 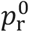 in session 1, for one participant in the 1-day (left) or 1-month (right) groups. Red: ‘enhance right’ group in session 2. Blue: ‘enhance left’ group in session 2. Line: the diagonal. **B**, Between-session correlation of 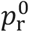 of sessions 1 and 3 for the two delay groups. Error bars: 95% confidence interval of the correlation, bootstrap. Magenta area: 95% confidence interval of the correlation expected under the assumption of complete stability, bootstrap Bernoulli processes. In contrast to the vertical bisection task, the correlation between 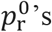’s in the two groups is comparable.

